# Hippocampal connectivity with sensorimotor cortex during volitional finger movements I. Laterality and relationship to motor learning

**DOI:** 10.1101/479451

**Authors:** Douglas D. Burman

## Abstract

Hippocampal interactions with the motor system are often assumed to reflect the role of memory in motor learning. Here, we examine hippocampal connectivity with sensorimotor cortex during two tasks requiring paced movements, one with a mnemonic component (sequence learning) and one without (repetitive tapping). Functional magnetic resonance imaging activity was recorded from thirteen right-handed subjects; connectivity was identified from sensorimotor cortex (SMC) correlations with psychophysiological interactions in hippocampal activity between motor and passive visual tasks. Finger movements in both motor tasks anticipated the timing of the metronome, reflecting cognitive control, yet evidence of motor learning was limited to the sequence learning task; nonetheless, hippocampal connectivity was observed during both tasks. Connectivity from corresponding regions in the left and right hippocampus overlapped extensively, with improved sensitivity resulting from their conjunctive (global) analysis. The cortical laterality of SMC connectivity depended both on the hippocampal source and the task.Functionally-defined seeds produced bilateral connectivity within the hand representation, regardless of whether finger movements were uni- or bimanual; these seeds were located midlateral within the hippocampus, whereas structural seeds were located in the posterior hippocampus and produced unilateral connectivity. Results implicate the hippocampus in volitional finger movements even in the absence of motor learning or recall.

## Introduction

Does the hippocampus play a role in executing volitional finger movements? There are several reasons to suspect it might. Although often assumed to reflect its known role in memory function (1–4), the hippocampus shows motor activity (5), especially during motor sequence learning (6–12). Furthermore, the hippocampus has been implicated in the generation of theta waves (13, 14), which enhances motor performance (15–17). Finally, low-threshold electrical stimulation of the hippocampus induces seizures (18), suggesting an intimate interaction with the motor system. These considerations suggest the hippocampus is intimately involved in movements, particularly volitional movements.

Paced movements require volitional movements, and the primary motor cortex is necessary for volitional movements of individual fingers (19, 20). Short fiber tracts connect postcentral with precentral regions (21), providing sensory feedback required for accurate performance. The psychophysiological interaction (PPI) technique for studying connectivity allows us to study task-specific hippocampal influences on SMC during these movements. In this technique, the hemodynamic signal is deconvolved to the underlying neural activity (22), then *interaction effects* between motor- and non-motor activity in the seed region (hippocampus) demonstrate differential effects on the magnitude or sign of response in the target region. This technique demonstrates the directional influence of connectivity on a moment-to-moment basis, important since the temporal pattern of hippocampal activity carries information (23–26).

This study explores the laterality and memory requirements of hippocampal-SMC connectivity during paced movements. Anticipatory motor responses confirmed a cognitive role during both sequence learning and paced, repetitive tapping. Hippocampal sources of connectivity were bilateral, whereas the laterality of SMC connectivity depended both on the task and seed; nonetheless, connectivity with the SMC hand representation increased during both tasks, suggesting connectivity was not limited to memory functions.

## Materials and methods

### Subjects

Thirteen right-handed adults from the Chicago metropolitan area participated in the study (ages 249, mean=42.3, five females). The nature of experimental procedures were explained to each subject before obtaining written consent; consent procedures complied with the Code of Ethics set forth in the Declaration of Helsinki, and were approved by the Institutional Review Board Board at the NorthShore University HealthSystem / Research Institute. Consented subjects had no history of the following: a previous concussion, psychiatric illness, learning disability, attention deficit disorder, abnormal physical or cognitive development that would affect educational performance, central neurological disease (or structural pathology), or neurosurgical intervention.

Immediately prior to their fMRI tests, subjects were trained on one cycle of each experimental task.

### Experimental tasks

During image acquisition, subjects performed a series of seven tasks, followed by five minutes of rest; these tasks were designed to reliably map diverse functions (27). The sequence of tasks was the same for all subjects, presented as follows: 2-back, visual association, visual/motor, Stroop, word association, and emotion. Each task consisted of a single run. Only results from the “visual/motor task” are reported here; results from the other tasks will be reported elsewhere.

The visual/motor task comprised 6 cycles of a specified sequence of visual and motor conditions. The timeline for a single cycle is diagrammed in Fig 1. Each cycle began with a 4s instruction screen, when a metronome ticked through sound-attenuated headphones [MR Confon Mk II] at 2 beats/sec as a 4-digit sequence appeared onscreen to specify the correct order of button presses. Once the digits were replaced with a cross, the subject pressed the remembered sequence of buttons synchronously with the metronome, repeating the 4-button sequence throughout the 16s block (8 repetitions). A visual block followed; a circular checkerboard pattern flickered at 4Hz for 9s, while subjects fixated the center of the pattern and refrained from moving.

**Fig 1.**
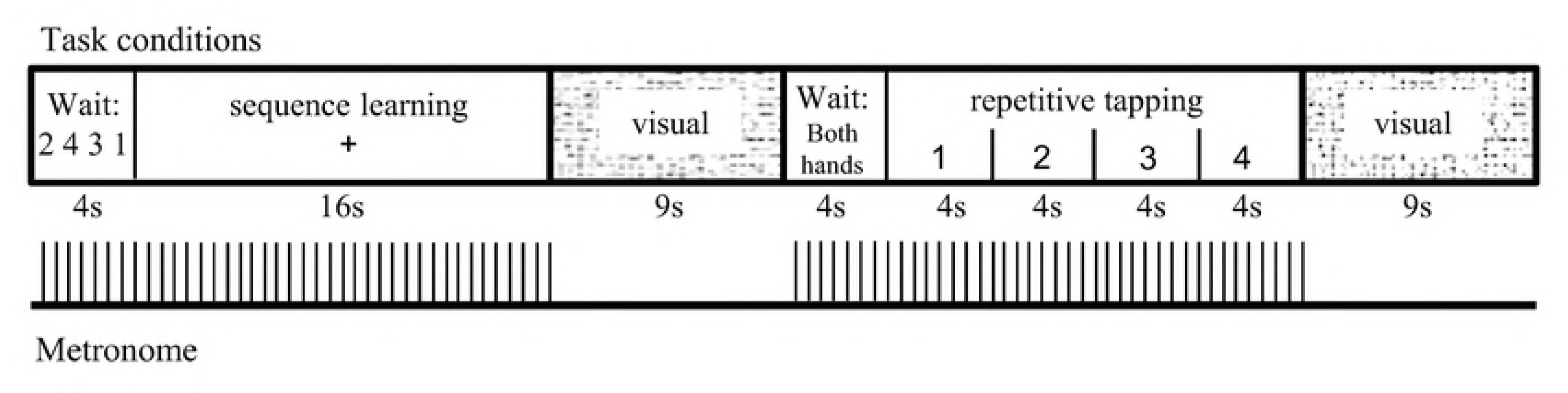
One cycle of the visual / motor task. Including instructions, two motor conditions alternated with a visual condition; this sequence repeated for a total of six cycles.

After the visual block, another 4s instruction screen instructed the subject to tap the same finger with both hands on cue. Once the instruction screen was replaced with the number ‘1’, the subject tapped the index finger from both hands in synchrony with the metronome; every 4s, the onscreen number increased by one, and the subject changed finger. Each finger tapped for 8 repetitions during the 16s repetitive tapping block; the cycle ended with another visual block.

With this design, each motor condition (with its instruction screen) was preceded and followed by a period of rest. Button presses during sequence learning were performed with the dominant (right) hand, with a new sequence specified at the beginning of the first 4 cycles; the first 2 sequences were repeated at the end to test for motor recall. Button presses were recorded from the right hand during both sequence learning and repetitive tapping.

### Behavioral analysis and terminology

Four buttons, arranged horizontally on the response pad, were each pressed by a different finger, labelled sequentially as F1 (index finger) through F4 (little finger). Movement of each finger was specified during the repetitive tapping blocks in 4s time periods, designated as T1 - T4 (reflecting movements of F1 - F4, respectively). The corresponding finger representations in sensorimotor cortex will be referred to as SM(F1) - SM(F4).

In the sequence learning block, a subject repeatedly tapped a 4-button sequence in synchrony with the metronome, beating at 500ms intervals. With four finger movements during each sequence, several behavioral measures were possible. In this study, three behavioral measures were calculated for each repetition: the mean stimulus-response asynchrony (the time interval between the metronome beat and the button press), the mean intertap interval, and the mean precision of intertap intervals (i.e., the absolute difference between 500ms and the intertap interval). Each mean value represented 3 or 4 measurements from a single sequence, so the standard deviation was calculated as a measure of the individual subject's response variability.

The intertap interval can be used as an example of our general method for visualizing and analyzing behavioral data. The mean intertap interval across all subjects was plotted (i.e., the group mean of the individuals’ mean intertap intervals), and a 2-tailed t-test evaluated whether the intertap intervals during the final two rehearsals significantly differed from the initial two rehearsals of the sequence. This test used *between-subject* variability to evaluate group changes in the mean intertap interval; if each subject initially had a mean intertap interval of 500ms, a consistent drop of 10ms in this mean group value would be significant, even if each subject had a standard deviation of 100ms.

The mean standard deviation across all subjects was also plotted (i.e., the group mean of the individuals' standard deviations). A 2-tailed t-test evaluated whether the variability in intertap intervals during the final two rehearsals differed from the initial two rehearsals. This test evaluated changes in *within-subject* variability; if each subject began with a standard deviation of 100 ms, consistent changes to 80 ms would be significant, even if the mean intertap interval for the group remained the same at 500 ms.

Stimulus-response asynchrony differentiated between reflexive movements (consistent short-latency responses following stimulus presentation) vs. intentional movements under cognitive control (anticipatory responses that precede the metronome sounds). Changes in intertap intervals and their precision identified learning effects resulting from rehearsal.

To compare behavioral performance between motor tasks, the same behavioral measures and statistical tests were additionally applied to 4-tap groupings during the repetitive tapping task. A 2-tailed t-test identified task differences in performance, both for the initial and final pair of rehearsals.

### MRI data acquisition

Images were acquired using a 12-channel head coil in a 3 Tesla Siemens scanner (Verio). Visual stimuli projected onto a screen (Avotec Silent Vision) were viewed via a mirror attached to the head coil, and behavioral responses were recorded by Eprime [Psychology Software Tools, Inc.] from an optical response box (Current Designs, Philadelphia, PA). Blood-oxygen level dependent (BOLD) functional images were acquired using the echo planar imaging (EPI) method, using the following parameters: time of echo (TE) = 25 ms, flip angle = 90°, matrix size = 64 x 64, field of view = 22 cm, slice thickness = 3 mm, number of slices = 32; time of repetition (TR) = 2000 ms. The number of repetitions for the visual/motor condition was 182. A structural T1 weighted 3D image (TR = 1600 ms, TE = 3.46 ms, flip angle = 90°, matrix size = 256 x 256, field of view = 22 cm, slice thickness = 1 mm, number of slices = 144) was acquired in the same orientation as the functional images.

### fMRI data processing

Data was analyzed using SPM8 software (http://www.fil.ion.ucl.ac.uk/spm). Images were spatially aligned to the first volume to correct for small movements; no run showed more than 4mm displacement along the x, y or z dimension. Sinc interpolation minimized timing-errors between slices; functional images were coregistered to the anatomical image, normalized to the standard T1 Montreal Neurological Institute (MNI) template, and resliced at 4mm^3^ resolution. Data were smoothed with a 10mm isotropic Gaussian kernel. Statistical analyses at the first level applied a high pass filter with a cutoff of 128s, applied to a design matrix with all 23 conditions from the 7 tasks. The instruction screen was treated as a covariate of no interest, and another covariate specified each transition between runs (28–30). Conditions of interest were specified for sequence learning, visual, and each of the four fingers moving during the repetitive tapping block; the latter four conditions were combined to identify brain activity across the entire repetitive tapping block. Blocks were modeled for analysis using a canonical hemodynamic response function (HRF).

Global normalization scaled the mean intensity of each brain volume to a common value to correct for whole brain differences over time. A parameter estimate of the BOLD response to each condition was generated; motor activation was identified by contrasting mean BOLD responses to motor vs. visual conditions. For group analysis, BOLD contrasts from individual subjects during the motor memory and repetitive tapping conditions were entered into a two-sample t-test. Analyses used an intensity threshold of p<0.05 with a family-wise error (FWE) correction for multiple comparisons, applied to a sensorimotor region of interest (ROI). This ROI was derived from the WFUPickatlas toolbox, and was specified as the overlap between TD and aal atlas labels for post-plus precentral gyrus.

This statistical model allowed us to identify activation by each motor condition, but also to identify activation in common through a global conjunction analysis of both conditions. Global analysis provided a sensitive measure for identifying areas activated in common by accounting for differences in variability for each condition; the resulting bilateral activation map for hand movements was used as the ROI for connectivity analysis.

### Laterality analysis

Laterality was assessed using the following formula:

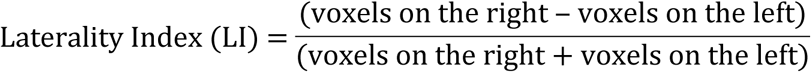

This provided a continuous variable, ranging from LI = −1 (activation or connectivity limited to the left hemisphere) to LI = +1 (activation or connectivity limited to the right hemisphere). Laterality was used to assess hemispheric differences in activation for a unilateral condition (sequence learning), hemispheric differences in activation for a bilateral condition (repetitive tapping), and hemispheric differences in sensorimotor connectivity during both conditions. These comparisons served to differentiate between SMC laterality differences attributable to unimanual vs. bimanual movements.

### Psychophysiological interactions (PPI)

#### Preprocessing

Connectivity analysis was carried out using psychophysiological interactions (22), modified to account for individual variability in connectivity (31). A total of 156 voxels were identified from the left (78) and right hippocampus (78) of the normalized brain, as delimited by the aal atlas in the WFU PickAtlas toolbox (http://fmri.wfubmc.edu/software/PickAtlas). Alternate voxels in the medial/lateral and anterior/posterior dimensions were sampled, beginning at the posterior/medial edge of the hippocampus. This grid of alternating voxels minimized artifactual effects from smoothing; it yielded a total of 13 voxels from the left and 13 from the right hippocampus. Coordinates for these voxels are specified in Table S1.

A contrast was selected and specified to create eigenvariates for all conditions in the statistical model at each hippocampal voxel (assured by selecting a p-value threshold of 1). An interaction term specified a greater effect of motor activity in the seed than during the visual condition; after adjustments for regional differences in timing and baseline activity, a regression analysis showed the magnitude of the BOLD signal that correlated with this interaction term elsewhere in the brain.

#### Seed selection

Appropriate selection of seed regions for connectivity analysis is critical (32), but because the temporal pattern of hippocampal activity affects information processing elsewhere (33–37), identifying areas of increased hippocampal activity through activation analysis may be inadequate. We therefore adopted two approaches. The first approach was structural; the hippocampus was divided into 8 regions from posterior to anterior, with connectivity evaluated for each region (position-1 through position-8). The second approach was empirical or functional; the hippocampal voxel generating the greatest connectivity anywhere within SMC during a task was selected as a seed.

A functional seed was identified from the left and right hippocampus for both the sequence learning task (functional seed-1) and the repetitive tapping task (functional seed-2). Connectivity from each functional seed (as well as structural seeds) was evaluated for both motor tasks.

#### Group analysis

Beta estimates of connectivity from each subject's left and right hippocampus were entered into a two-sample t-test, using the hand representation as the ROI and an intensity threshold of p<0.05 with FWE correction. This ROI reflected our primary hypothesis: hippocampal connectivity with the finger region of SMC increases when initiating volitional finger movements. In addition to connectivity maps from each hippocampus, global analysis of connectivity from both hippocampi identified joint effects, after accounting for hemispheric variability in connectivity.

#### Individual analysis

Individual results were examined to ensure that group results were representative, both regarding laterality and the overlap in connectivity between seeds from the left and right hippocampus. Thresholds for these displays were set at 50% of the maximal connectivity value, providing connectivity maps comparable in extent to those observed from group analysis.

#### Testing for Cortical Selectivity

Two approaches were used to test for cortical selectivity. First, global analysis of connectivity was tested individually in both hand and foot representations, followed by a 1-way ANOVA to directly compare the amplitude of connectivity between areas. (The foot representation was identified from activation in a different cohort.) Second, the hypergeometric distribution (38) calculated the probability that the observed number of connectivity voxels reaching statistical significance overlapped the hand representation in the left or right SMC by chance. Cortical selectivity was demonstrated by a probability <5% (p<0.05).

The hypergeometric distribution compared the number of voxels reaching criterion in-vs. outside the hand representation; the same t-value threshold was applied to each, using different ROIs (the SMC in-vs. outside the hand representation). This approach was adopted due to the method for selecting functional seeds, which identified the single highest connectivity value for a subject anywhere within SMC. Data review indicated that no two subjects showed maximal connectivity at the same voxel, so significant connectivity for the group of subjects was not assured anywhere within SMC -- but if observed, what is the likelihood it would be restricted to the hand representation? In other words, given that connectivity values meeting a statistical threshold could hypothetically appear anywhere within SMC, what is the probability that the actual number of voxels meeting this threshold within the hand representation would occur by chance? Analogous to drawing white and red balls from an urn without replacement, the probability of voxels meeting threshold *inside* the hand representation by chance (drawing *n* white balls) out of all the voxels in SMC (a total of *m* balls) depends on the total number of SMC voxels in- and outside the hand representation (the total number of white and red balls).

This approach assumes the connectivity threshold could be exceeded anywhere within SMC, i.e., the interaction term between motor and visual conditions in the hippocampus does not merely reproduce the activation analysis used to identify the SMC hand representation. In fact, these methods differ markedly. Activation analysis identified where the mean SMC activity during motor blocks exceeded the mean activity during visual blocks; by contrast, the interaction term used in PPI analysis reflected the temporal pattern of hippocampal activity throughout both blocks.

#### Independence of activation and connectivity

Fig 2 shows differences between activation and PPI connectivity, illustrating their independence. During activation analysis, a motor condition (sequence learning or repetitive tapping) was contrasted with the passive visual condition; based upon the block design, the predicted BOLD response is the same across all 6 task cycles (Fig 2A, blue waveform at top). For each task, the predicted signal was elevated above baseline during the motor block (shaded gray), and depressed below baseline during the visual condition. By contrast, the BOLD response predicted from connectivity analysis depends on the pattern of seed activity throughout the task. The green and red traces at the bottom of Fig 2A show one subject’s predicted BOLD response based on connectivity, derived from activity in the left and right hippocampal seeds, respectively. Note that the predicted pattern of activity changes across cycles, and that predicted increases in the BOLD response above baseline often do not coincide with those predicted from activation analysis. Indeed, BOLD responses predicted from the left and right hippocampal seeds show greater similarity to each other than to the response predicted from activation.

**Fig 2.**
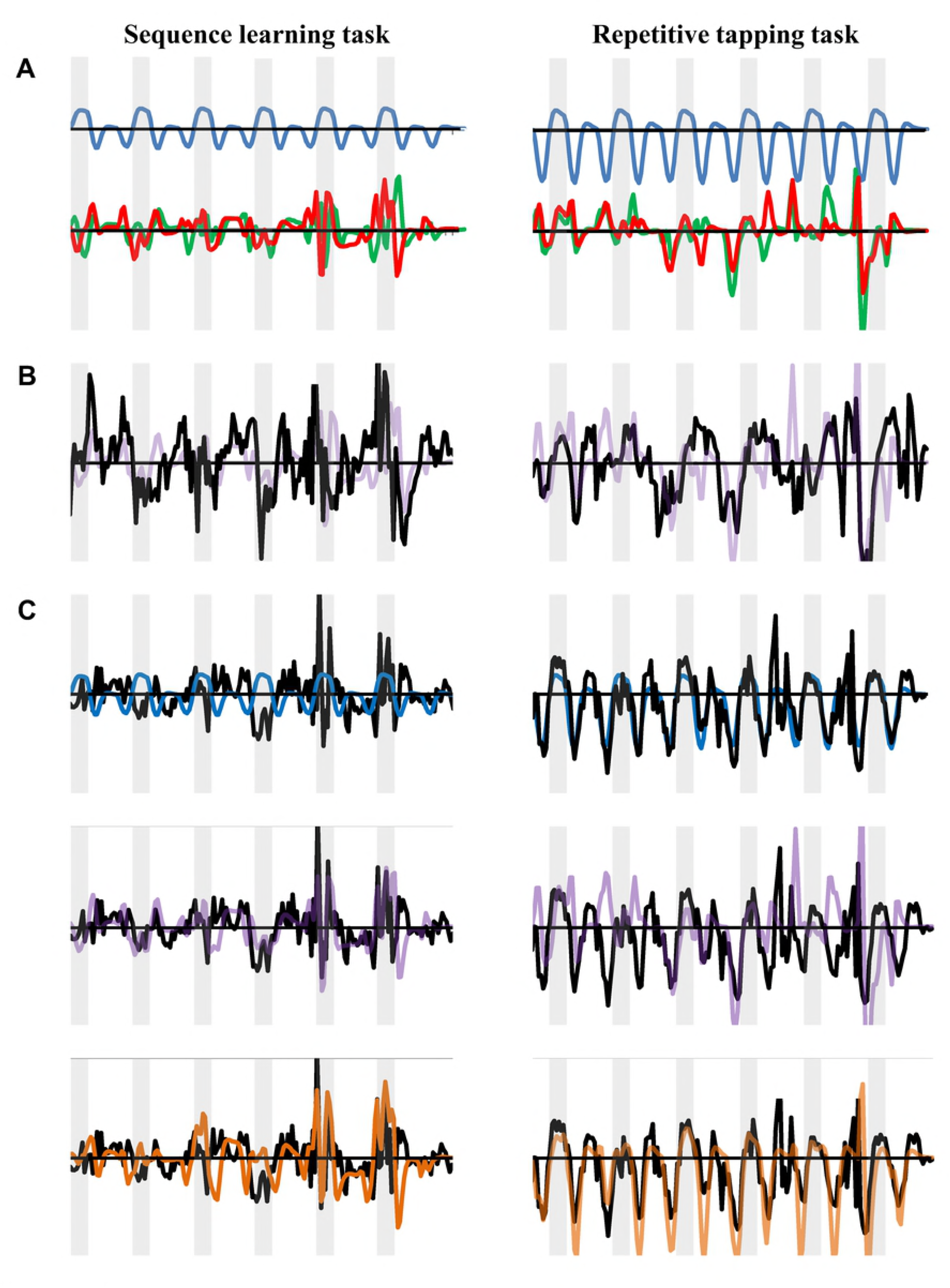
Independence of activation and connectivity analysis. (A) Predicted patterns of BOLD activation (top, blue), and connectivity from the left and right hippocampal seeds (bottom, green and red, respectively). (B) BOLD activity observed at the SMC connectivity maxima (black) compared with activity predicted from the sum of the activity from the left and right hippocampal seeds (purple). (C) BOLD activity observed at the SMC activation maxima (black) compared to the predicted pattern for activation (top, blue), the sum of activity from left and right hippocampal seeds (middle, purple), and the sum of activation plus hippocampal activity (bottom, orange).

The BOLD signal observed at the SMC *connectivity maxima* (Fig 2B, black line) approximated the sum of BOLD signals predicted from combined activity in the left and right seeds (purple line). In the sequence learning task (left), the peaks and valleys at this right SMC voxel generally did not coincide with those predicted from activation analysis; although connectivity was significant at this voxel, activation was not. In the repetitive tapping task (right), the BOLD signal at the connectivity maxima again approximated the sum of BOLD signals predicted from activity in the left and right seeds; due to sufficient overlap with the pattern predicted from the task design, there was also significant activation.

For both tasks, the BOLD signal at the *activation maxima* approximated the response predicted from the task design (Fig 2C, top, blue line), but also the response predicted from connectivity analysis (Fig 2C, middle, purple line). However, the BOLD signal most closely mirrored the response predicted from the sum of activation and connectivity analyses (Fig 2C, bottom, orange line).

Thus, the pattern of BOLD activity in a subject’s SMC could reflect the pattern of seed activity predicted from connectivity analysis, activation analysis, or both.

## Results

### Behavioral performance during motor tasks

Stimulus-response asynchrony, intertap intervals, and precision of intervals were characterized for both sequence learning and repetitive tapping tasks (Fig 3, see also Tables 1 and 2). The left column shows group changes in performance across rehearsals, calculated from mean values for all subjects; the right column shows the mean standard deviation among individuals.

**TABLE 1.**
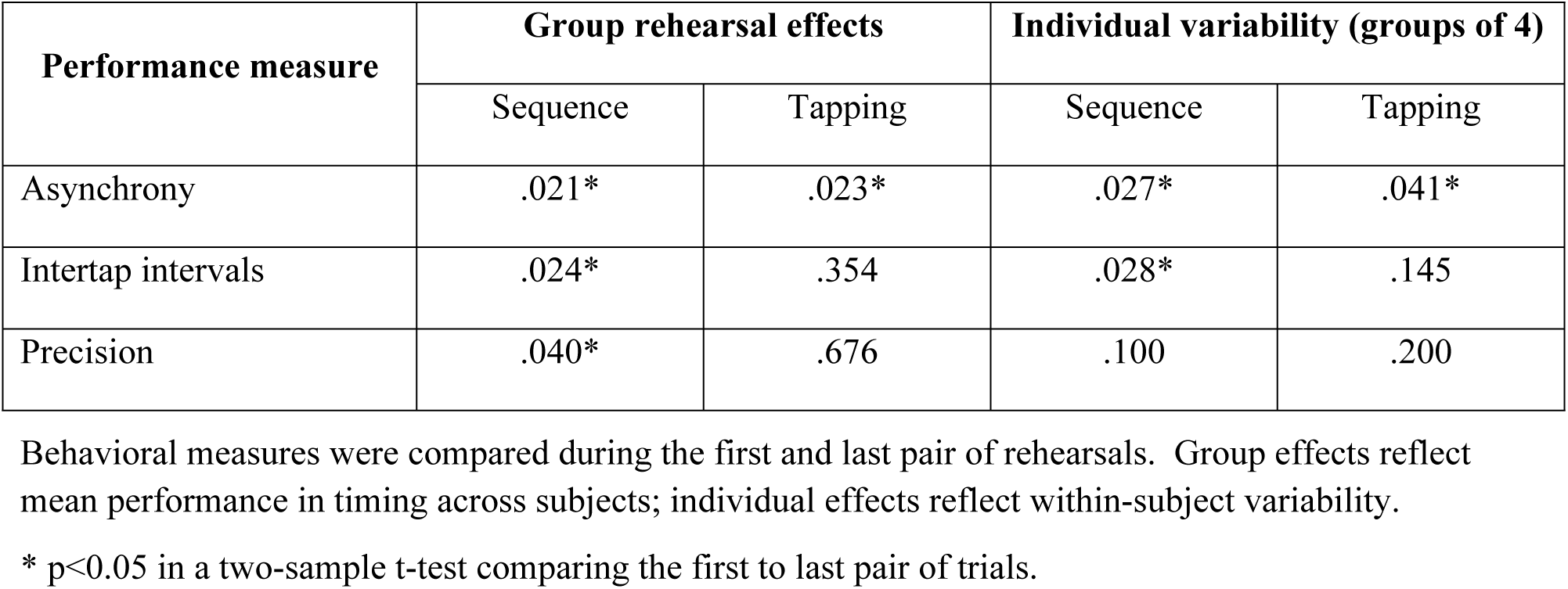
Rehearsal effects in sequence learning and repetitive tapping tasks.

**TABLE 2.** Task differences during performance of sequence learning and repetitive tapping tasks.

| Performance measure | Group performance effects |  | Individual variability (groups of 4) |  |
| --- | --- | --- | --- | --- |
|  | Trials 1-2 | Trials 7-8 | Trials 1-2 | Trials 7-8 |
| Asynchrony: |  |  |  |  |
| 1st Sequence vs. Tap | .500 | .589 | .006* | .103 |
| Repeat Sequence vs. Tap | .600 | .013* | .130 | .963 |
| 1st Sequence vs. Repeat | .101 | .008* | .145 | .176 |
| Intertap intervals | .687 | .531 | .046* | .102 |
| Precision | .782 | <.001* | .425 | .825 |

Task differences were identified from initial performance and again following rehearsal. Performance following presentation of a novel sequence in the learning task was distinguished from its repeat presentation.
\* $p < 0.05$ in a two-sample t-test.

**Fig 3.**
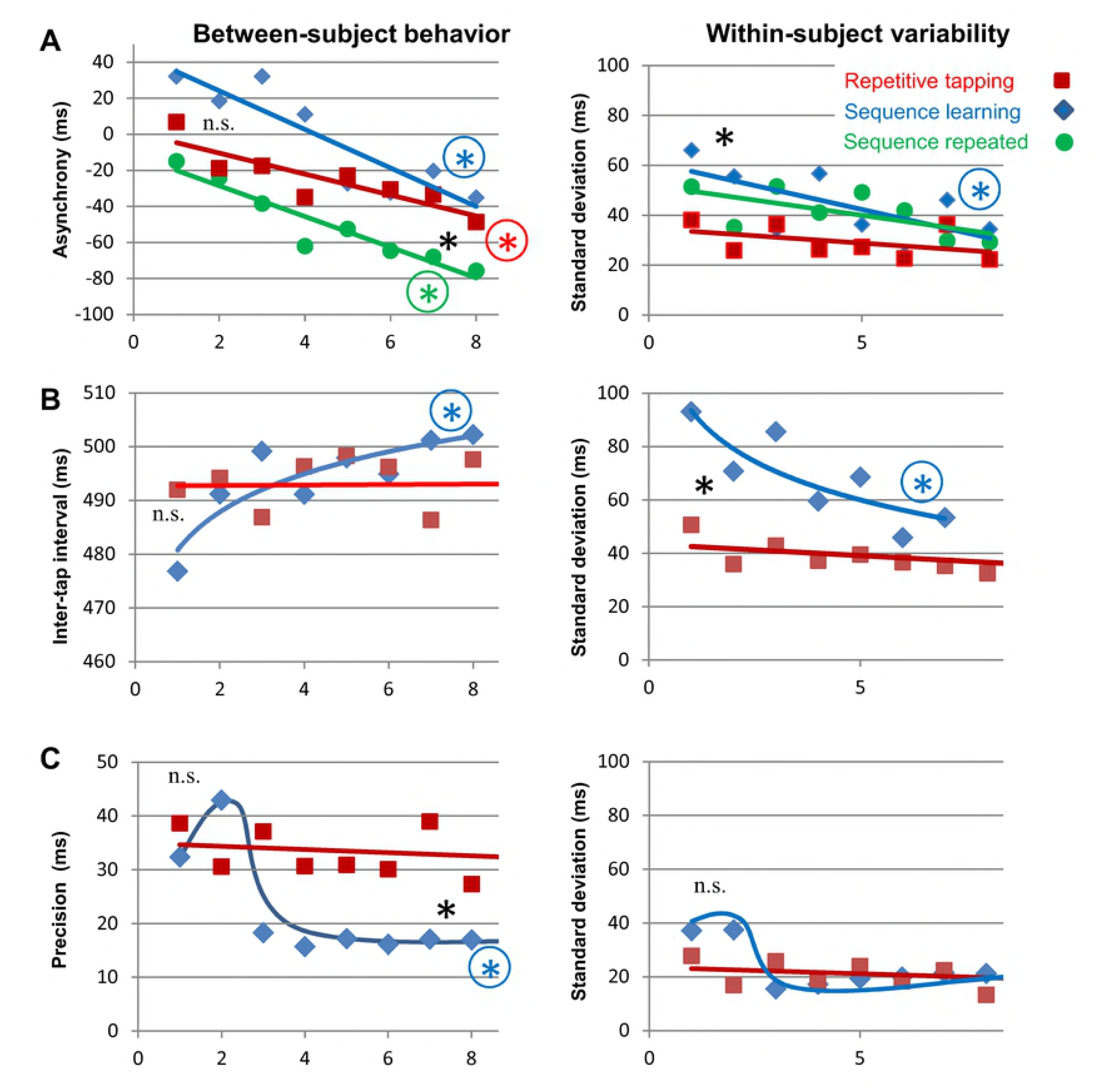
Rehearsal effects on motor performance during sequence learning and repetitive tapping. (A) Changes in stimulus-response asynchrony across rehearsals. Negative values for asynchrony represent anticipatory responses, which developed across rehearsals for all motor conditions; variability represents standard deviations for 4-note groupings among individuals. Colored encircled asterisks indicate a significant difference from the first to last pair of repetitions during a task; a plain black asterisk at the beginning or end of rehearsals indicates a significant difference between task conditions (p<0.05). (B) A change in intertap intervals was observed across rehearsals only in the sequence learning task, accompanied by a decrease in variability. (C) A change in precision was observed across rehearsals only in the sequence learning task.

Fig 3A shows the effects of rehearsal on stimulus asynchrony (i.e., the time interval between the metronome and button presses). The desired pacing of finger tapping was set by a metronome during the instruction period. Finger movements during the movement task began synchronous or slightly after the metronome (left), but significantly anticipated the metronome with rehearsal (negative values). Repetitive tapping (red) and sequence learning (blue) generated similar anticipatory responses after 8 rehearsals, but an additional 8 rehearsals on a learned sequence generated greater asynchrony (green). Individual *variability* in asynchrony was initially greater during sequence learning, but differences were eliminated over 8 rehearsals as performance of a learned sequence became easier (right).

Learning effects were identified from rehearsal-related changes in intertap intervals and their precision (Fig 3B-C). Learning effects were observed during sequence learning but not the repetitive tapping task; the intertap interval during sequence learning significantly improved towards the paced interval of 500 ms, while its variability dropped. The precision of intertap intervals also improved during sequence learning (i.e., how much intertap intervals within a 4-key sequence differed from 500 ms), dropping below that observed during repetitive tapping.

### SMC activation during motor tasks

SMC was activated during both sequence learning and repetitive tapping (Fig 4a and Table 3). Reflecting unimanual performance by the right hand, SMC activation during sequence learning was limited to the left hemisphere during group analysis (LI = −0.98), but not during individual analysis (mean LI = −0.47±0.33). By contrast, group activation during repetitive tapping was right-dominant (LI=0.62), whereas activation during individual activation showed equal dominance (mean LI = 0.08±0.41), reflecting bimanual performance. Thus, group analysis of individual tasks poorly reflected the extent of motor activity present among individuals.

A more sensitive method for identifying the extent of motor activity during group analysis involved global (conjunctive) analysis of both motor conditions (Fig 4B). Global analysis extended the area of group activation (cyan, see especially the right SMC for sequence learning), while overlapping most activation observed during single-task analysis (orange). Whereas global analysis of *group* activation better reflected the full extent of activation observed during individual analyses, global analysis of *individual* activation reduced the apparent area of activation.

**TABLE 3.** Location of group sensorimotor cortical activation.

| Task | Hemisphere | Cluster size | Z-score<br>(peak) | Peak Coordinates |
| --- | --- | --- | --- | --- |
| Sequence learning | left | 227 | 6.08 | (-42,-36,54) |
|  |  |  | 5.69 | (-42,-4,46) |
|  |  |  | 5.67 | (-38,-8,54) |
| Repetitive tapping | right | 157 | 5.04 | (42,-20,54) |
|  |  |  | 5.01 | (46,-16,50) |
|  | left | 71 | 4.69 | (-42,-4,46) |
|  |  |  | 4.27 | (-50,0,38) |
| Combined<br>(global analysis) | left | 165 | 5.82 | (-46,-36,54) |
|  |  |  | 5.65 | (-42,-4,46) |
|  |  |  | 5.07 | (-50,0,38) |
|  | right | 168 | 5.16 | (46,-32,50) |
|  |  |  | 5.05 | (46,-16,50) |
|  |  |  | 5.00 | (42,-24,54) |

Activation clusters within the sensorimotor mask, surviving an intensity threshold of $p=0.05$ with a family-wise error correction and extent threshold of 20. Peak locations are specified in MNI coordinates.

**Fig 4.**
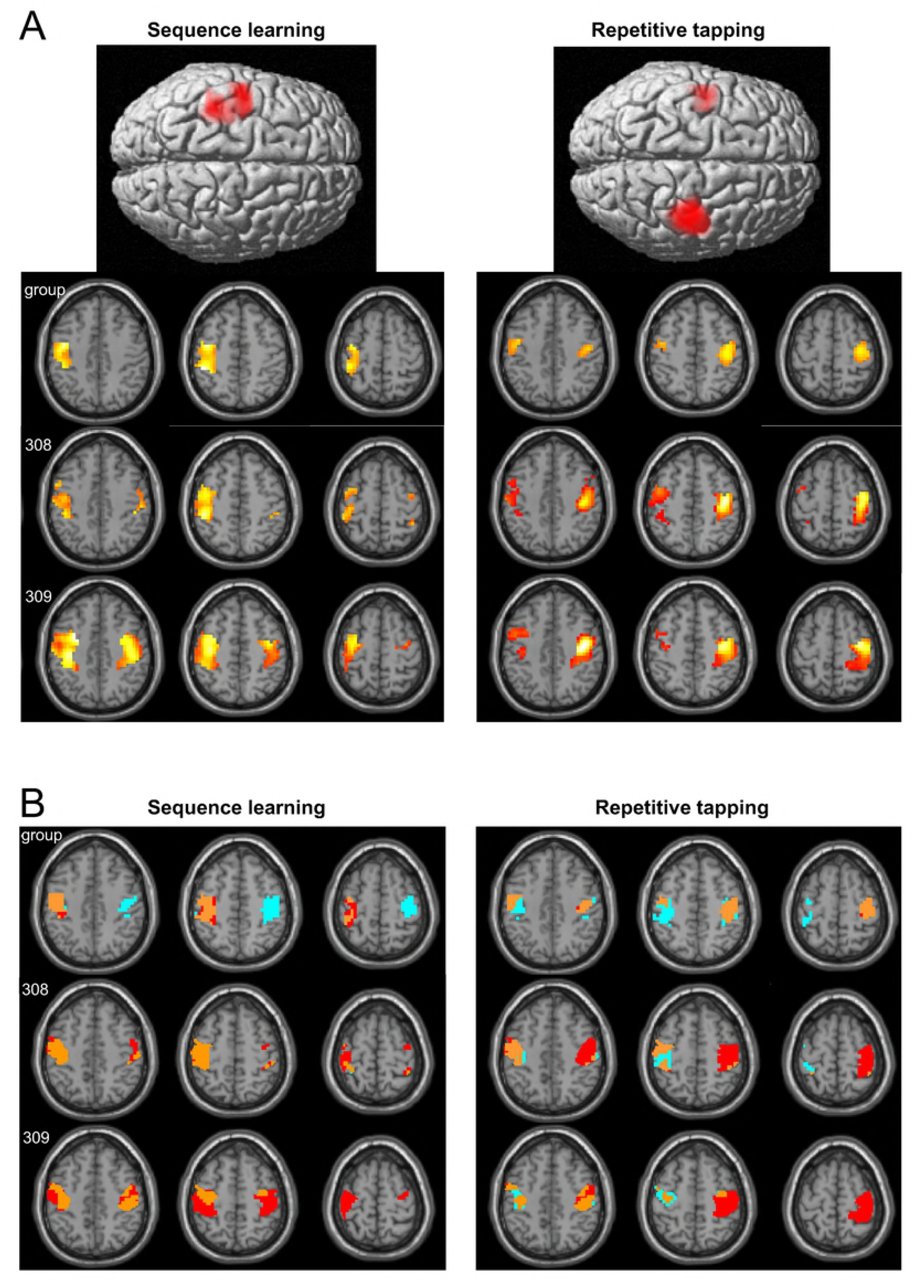
Sensorimotor activation during performance of motor tasks. (A) Group analysis (top) showed unilateral activation in left sensorimotor cortex during performance of the unimanual sequence learning task, and bilateral activation during the bimanual repetitive tapping task. Individual analysis (bottom) showed bilateral activation for both tasks. (B) Global conjunctive analysis of both motor conditions demonstrated bilateral activation during group analysis (top) as well as individual analysis (bottom); global analysis for the group expanded the area of demonstrable activation. Activation observed exclusively during individual task analysis is shown in red, exclusively during global analysis in cyan, and overlapping activation in orange. Thresh old for activation was p=0.05 withF"WE correction for multiple comparisons. Results in this and subsequent figures are displayed with the neurol ogi cal conventi on (left di splay = 1 eft side of brain).

In Fig 4, activation was explored using the SMC mask in order to identify the full hand representation. Global conjunctive analysis without a mask demonstrated additional activation in the supplementary motor cortex and cerebellum, plus auditory activation from the metronome in the superior temporal gyrus (Table S2).

### Hippocampal sensorimotor connectivity

Hippocampal connectivity was evaluated within the SMC hand representation. Although no significant differences in connectivity were observed between tasks, regardless of seed source, the pattern of results during group analysis depended both on the source of the seed and the task (Fig 5A and Table 4). For functional seeds, connectivity from either the left (red) or right hippocampus (yellow) typically generated right-dominant connectivity, with extensive overlap in connectivity from the left- and right-hippocampus. Overlap from left and right seeds invariably exceeded 50%, with a mean overlap of 85.0±32% for repetitive tapping and 86.5±15% for sequence learning. Global analysis of connectivity from both hippocampi demonstrated additional connectivity (cyan), particularly in the left SMC during sequence learning.

**TABLE 4.** Connectivity clusters within sensorimotor cortex during group analysis.

| Task | Seeds | Hemisphere (SMC) | Region(s) | Cluster size | Z-score (peak) | Peak Coordinates |
| --- | --- | --- | --- | --- | --- | --- |
| Sequence learning | Funct-1 | right | pre/post | 138 | 6.00<br>5.55<br>5.35 | (46,-24,58)<br>(38,-12,66)<br>(38,-32,62) |
|  |  | left | pre/post | 49 | 3.77<br>3.71 | (-30,-24,58)<br>(-38,-36,62) |
|  | Funct-2 | right | pre/post | 121 | 4.71<br>4.63<br>4.25 | (42,-28,62)<br>(38,-12,66)<br>(46,-8,58) |
|  |  | left | pre/post | 137 | 4.16<br>3.89<br>3.50 | (-38,-36,62)<br>(-34,-20,58)<br>(-50,-16,46) |
|  | Struct-1 | left | post | 46 | 4.82<br>3.49 | (-42,-36,58)<br>(-34,-20,46) |
|  | Struct-2 | -- | -- | -- | -- | -- |
| Repetitive tapping | Funct-1 | left | pre/post | 47 | 3.66 | (-34,-20,58) |
|  |  | right | pre | 8 | 3.57 | (34,-12,62) |
|  | Funct-2 | right | pre/post | 166 | 5.19<br>4.95<br>4.41 | (34,-16,58)<br>(46,-24,54)<br>(38,-32,58) |
|  |  | left | pre/post | 123 | 4.03<br>4.02<br>3.71 | (-38,-36,62)<br>(-34,-20,58)<br>(-50,-24,46) |
|  | Struct-1 | right | pre/post | 127 | 3.94<br>3.88 | (38,-20,50)<br>(34,-20,62) |
|  |  |  |  |  | 3.75 | (38,-12,66) |
|  |  | left | pre | 3 | 3.06 | (-30,-20,54) |
|  | Struct-2 | right | pre/post | 39 | 3.31 | (38,-23,66) |
|  |  | left | pre | 3 | 3.11 | (-34,-24,54) |

**Fig 5.**
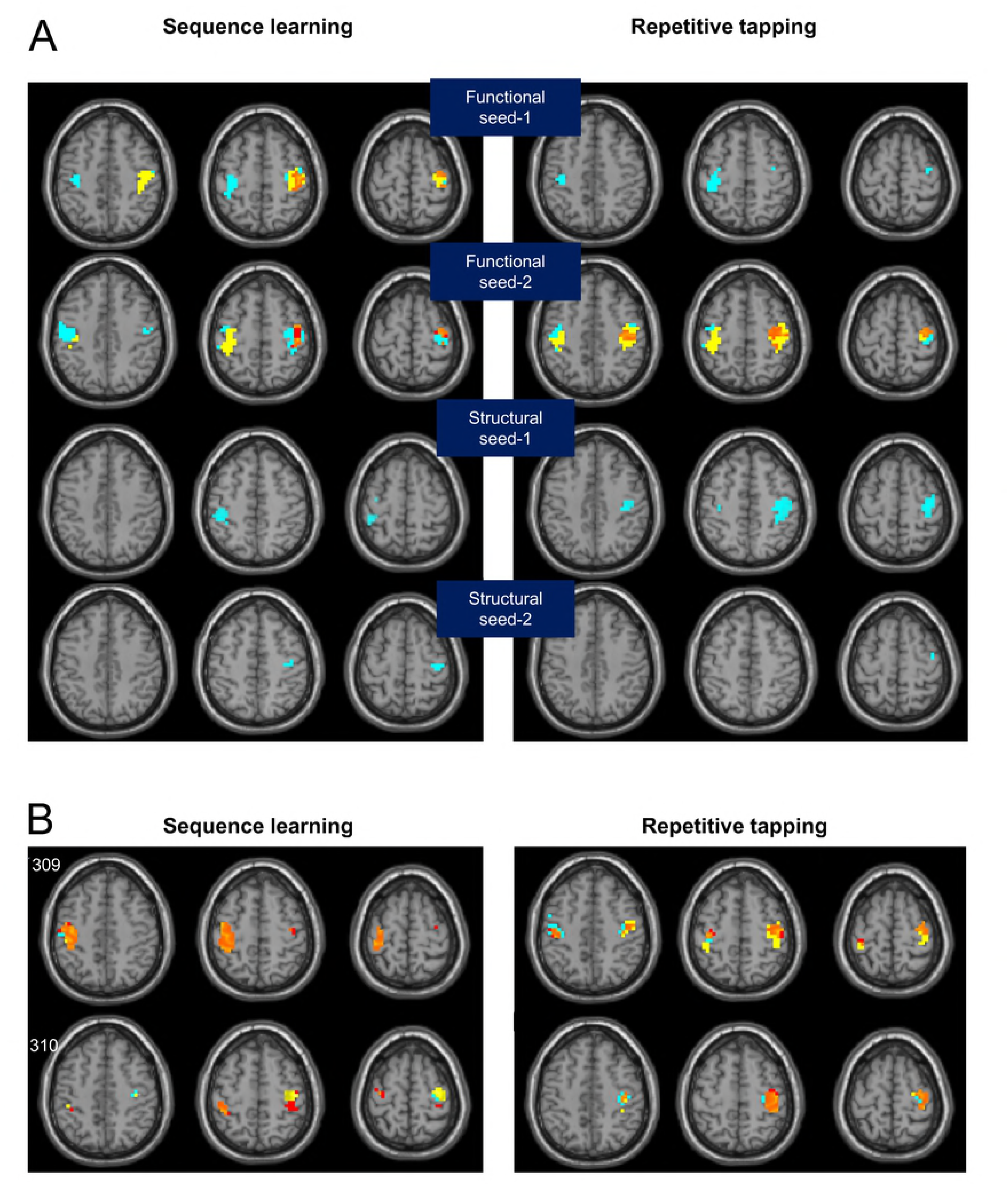
Sensorimotor connectivity from the left and right hippocampus during motor performance. (A) The laterality of connectivity depended on both the seed and motor task. For functionally-defined seeds, connectivity from the left (red) and right hippocampus (yellow) overlapped extensively in the right hemisphere (orange); additional regions of connectivity were evident from global analysis of both (cyan). (B) Individual analysis of sensorimotor connectivity from functional seeds; individual connectivity values exceeding 50% of maximum are displayed.

Connectivity from structural seeds was only observed from global analysis, but the pattern of connectivity again depended on the task and seed. Connectivity from structural seed-1 was left dominant during the sequence learning task but right dominant during repetitive tapping; connectivity from structural seed-2 was right dominant for both tasks, although more extensive during the sequence learning task.

Individual analysis showed extensive overlap from the left and right hippocampal seeds in both hemispheres (Fig 5B), with bilateral connectivity during sequence learning despite unimanual task performance.

### Cortical selectivity for hand representation

During ROI analysis, neither functional seed-1 nor functional seed-2 generated significant connectivity within the SMC foot representation.

Cortical selectivity for the hand representation was also evaluated with the hypergeometric distribution; this identified the chance probability of the observed number of voxels showing connectivity within the hand representation, relative to the total number of voxels showing connectivity throughout SMC (Table 5). Functional seeds showed significant selectivity for the hand representation in both hemispheres, except the left hemisphere for functional seed-2 during the repetitive tapping task. Due to widespread connectivity elsewhere in SMC, selectivity was not significant in this task, despite connectivity for 123 out of 165 voxels within the hand representation.

**TABLE 5.**
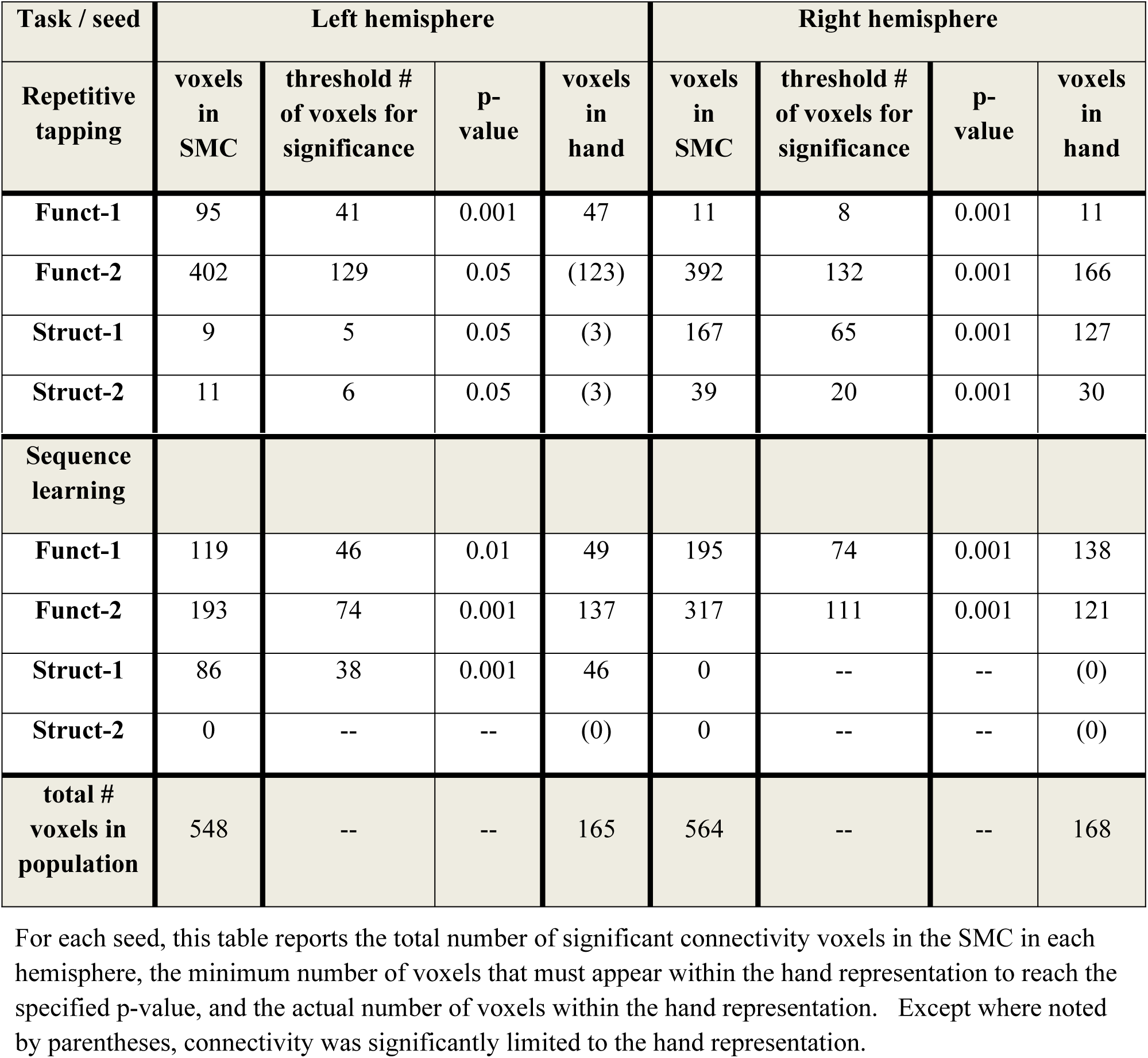
Statistical significance of overlap between motor-related connectivity with the hand representation using the hypergeometric distribution.

### Hippocampal source of motor connectivity

Because different regions of the hippocampus vary in their functional properties and anatomical connections (39–43), voxels functionally involved in sensorimotor control should aggregate within a restricted region of the hippocampus. Voxels associated with functional seeds aggregated near the lateral edge (see Fig S1; chi squared = 61.020, p < 0.001), whereas structural seeds were posterior.

## Discussion

Connectivity of the hippocampus with sensorimotor cortex (SMC) was examined during two paced motor tasks; behavioral analysis showed that performance during both tasks was under volitional control, although only one task produced learning effects. In both tasks, contributions from the left and right hippocampus overlapped, jointly generating bilateral connectivity in SMC; this connectivity was limited to the hand representation, identified from task activation across both tasks. These findings demonstrate a specific hippocampal influence on SMC during volitional movements that is independent of its role in motor learning.

### Behavioral analysis

Analysis of behavior during motor performance addressed the issue of volitional control and whether motor learning occurred across rehearsals.

Instructed to press a key in synchrony with the metronome ticks, presented at 500ms intervals, subjects consistently anticipated the metronome in both tasks. Reflecting subjects’ cognitive intent to tap at set intervals, the effect was additive across trials; if responses had been stimulus-driven, button presses would instead have followed each metronome tick at a fixed interval. Anticipatory responses thus reflected volitional control during both motor tasks.

Learning effects were evident during sequence learning, including improved precision in intertap intervals and decreased variability across rehearsals, whereas no such effects were evident during repetitive tapping. Learning effects in the sequence learning task occurred over the first 3-4 rehearsals, when cognitive processes are important for skill acquisition (44).

### Sensorimotor activation

Consistent with previous reports, group analysis showed SMC activation within the omega spur contralateral to the moving hand(s) (45–49).

Bilateral SMC activation was generated by the bimanual repetitive tapping task. Sequence learning, performed with the right hand only, generated left SMC activation during group analysis, but bilateral activation during individual analysis. This bilateral activation likely reflects inhibitory influences between motor regions in opposite hemispheres (50). Because movements of individual finger and their representations are not independent (51, 52), however,activation must reflect some combination of excitatory and inhibitory processes in both hemispheres.

Global conjunctive analysis included both sequence learning and repetitive tapping, improving sensitivity to common motor effects by accounting for task differences in variability. Global analysis showed bilateral activation, used to identify the SMC hand representation.

### Hippocampal seeds as sources of sensorimotor connectivity

Both functional and structural seeds were used to identify hippocampal connectivity with SMC. The connectivity of functional and structural seeds-1 both showed differences in laterality between motor tasks, perhaps due to uni-vs. bimanual performance during sequence learning vs. repetitive tapping. Nonetheless, all seeds generating significant SMC connectivity did so for both motor tasks, indicating hippocampal modulation of sensorimotor cortical activity did not require motor learning.

Functional seeds were identified empirically from the region with the highest connectivity to SMC; they were localized laterally, mostly within the middle third of the hippocampus, whereas structural seeds were posterior. This pattern suggests functional specialization within the hippocampus. Such specialization is also suggested by different patterns of laterality observed across tasks by different seeds. While our study could not adequately evaluate the source of such functional differences, they might regulate diverse processes involved in generating even simple movements: excitatory and inhibitory influences between fingers on the same hand, coordination of flexor or extensor muscle movements, and inhibitory influences between motor regions in opposite hemispheres (50–52)

The left and right hippocampus show similar connectivity patterns along its long axis (53), despite laterality differences in hippocampal function (54, 55). For each of our seeds, connectivity from the left and right hippocampus overlapped, such that increased sensitivity was provided by global analysis of both. (Indeed, connectivity from structural seeds was only evident from global analysis.) Thus, similar hippocampal regions in the two hemispheres effectively worked together.

### Hippocampal sensorimotor connectivity

Although hippocampal connectivity with the striatum has been observed during motor learning, suggesting a mnemonic-motor interaction (Fernandez-Seara et al., 2009; Albouy et al., 2013; Albouy et al., 2015), this study is the first to find hippocampal interactions with the motor system independent of its role in memory. The demonstration of connectivity with the sensorimotor cortex implicates the hippocampus in volitional movements, rather than just striatal-associated movements derived from habits (19, 20, 56, 57).

The hippocampus is the likely source of theta EEG rhythms (Ekstrom et al., 2005; Pignatelli et al., 2012), which reflect cognitive conditions conducive to motor performance, learning, working memory, sensorimotor integration, or spatial learning (Axmacher et al., 2010; Kober and Neuper, 2011; Cruikshank et al., 2012; Lega et al., 2012). By identifying where SMC activity was influenced more by hippocampal activity during movements than during the passive visual condition, our PPI method of analysis was task-specific for motor function. Furthermore, hippocampal connectivity with SMC was limited to the bilateral hand representation which initiated the task-related movement.

### Cortical selectivity: The hand representation

PPI analysis assured motor specificity by identifying the interaction term, whereby the moment-to-moment hippocampal activity during movements showed greater influence on SMC activity than the passive visual condition. The magnitude of connectivity was based on the correlation of SMC activity with this movement-specific term.

Increased hippocampal connectivity was observed during both tasks, invariably restricted to the SMC hand representation in at least one hemisphere. The Funct-2 seed met our statistical test for cortical selectivity in the right but not left hemisphere during repetitive tapping, but even in the left hemisphere, the hand representation showed extensive connectivity (see Table 5, where 123 of 165 voxels in the hand representation met threshold). When ten or more SMC voxels in a hemisphere met our group threshold, connectivity from structural and functional seeds was otherwise selectively restricted to the hand representation.

This cortical specificity indicates the connectivity does *not* represent a widespread influence on sensorimotor cortex, which might have been expected, based on studies that implicate the hippocampus with theta waves conducive to motor performance (13–17).

### Functional implications

Known hippocampal functions related to memory and sensorimotor integration might partially explain the results of this study.

Hippocampal connectivity with SMC was observed during both sequence learning and repetitive tapping tasks, even though the latter task did not show motor learning effects. Memory for the 500ms pacing interval was still evident, however, as anticipatory movements preceded the metronome during both tasks; because the hippocampus responds differentially to the intervals between stimuli, at least within a sequence (58–60), hippocampal connectivity may have relayed its “memory” for temporal intervals. Note that subjects could have simply responded to the metronome, however, so task performance did not *require* memory for the pacing intervals, leaving open the question of how and why hippocampal connectivity was recruited.

The hippocampus has also been implicated in sensorimotor integration (Caplan et al., 2003; Ekstrom et al., 2005; Ekstrom and Watrous, 2014). During sensorimotor integration, motor responses to sensory stimuli are modified, such as a navigational response to sensory surroundings. In the current study, movements were not constrained by acoustic stimuli, evident from the anticipatory responses noted earlier; however, hippocampal connectivity might have modified somatosensory processing in the postcentral gyrus, promoting the appropriate finger movements. If so, this would likely require cognitive control for initiating movements, as short fiber tracts connect postcentral with precentral regions of sensorimotor cortex for sensory feedback (21).

Part II of this study further investigates the role of hippocampal connectivity in volitional movements by comparing properties of hippocampal connectivity with objective criteria for cognitive control.

## Conclusions

Hippocampus connectivity with the SMC hand representation was evident during volitional, paced finger movements; this connectivity did not require motor learning, but was selective for movements. Lateral and posterior regions of the hippocampus differed in laterality across tasks,suggesting functional specialization within the hippocampus, yet connectivity from corresponding regions of the left and right hippocampus overlapped extensively. The role of this connectivity appears to be linked, directly or indirectly, to the control of volitional movements.

## Acknowledgments

The author wishes to thank Donald J. Bolger for his suggestions on an early draft of this manuscript, and the Center for Advanced Imaging (CAI) at NorthShore University HealthSystem for its administrative support.

---

**Fig S1. Distribution of functional seeds generating sensorimotor connectivity during motor tasks**. Functional seeds were located lateral, extending from middle to anterior hippocampus.

